# Low-Cost In-House Re-formulated Brain Heart Infusion Medium for Effective Planktonic Growth and Early Detection of Bloodstream Bacterial Pathogens

**DOI:** 10.1101/2025.07.08.663789

**Authors:** Jonathan Hira, Nasib Bin Mahbub, Jawad Ali, Rafi Ahmad

## Abstract

Sepsis, a clinically defined life-threatening condition, is a global contributor to high morbidity and mortality rates in humans. It is caused by systemic bloodstream bacterial infections, primarily involving aerobic pathogens such as *Escherichia coli, Staphylococcus aureus*, and *Klebsiella pneumoniae*. Rapid and accurate identification of these pathogens is a high-demand task, as prolonged diagnosis may increase the mortality rate among sepsis patients. Worldwide, commercial blood culture systems, such as BD BACTEC™ PLUS Aerobic/F /F culture bottles (used in this study), are routinely used to monitor bloodstream infections. However, due to high costs ($10.00-$15.00/bottle), limited availability of culture media (especially in low- and middle-income countries, and war zones), and a lack of customization for antibiotic susceptibility assay and epidemiology research, there is a need for secondary alternatives to facilitate the growth and identification of bloodborne pathogens. Therefore, we developed a low-cost ($4–$5/bottle) in-house culture medium with a newly improved formulation of Brain Heart Infusion media that enhances bacterial growth from spiked human blood tested on a panel of bacteria (*Escherichia coli, Staphylococcus aureus, Klebsiella pneumoniae, Acinetobacter baumannii, Pseudomonas aeruginosa*, and *Enterococcus faecalis*). The growth dynamics of these microbes in in-house formulated BHI-Blood+ culture media coincide with those in BACTEC™ Plus Aerobic/F culture vials, which primarily suggests the compatibility of bloodborne pathogens with this media and can be flagged positive <8h based on cellular growth rate. Additionally, conventional qPCR-based early detection (< 24h) and validation with the Oxford Nanopore MinION NGS platform highlight the value of this in-house culture media as an alternative to commercial culture media in terms of low-cost availability.

## Introduction

Bloodstream infections (BSIs) are severe conditions caused by the presence of viable pathogens, including bacteria, fungi, or viruses, in the bloodstream, often progressing to sepsis, a life-threatening condition. Sepsis arises when localized infections, such as those in organs, trigger a dysregulated immune response, leading to systemic inflammation, organ dysfunction, and potentially septic shock [1]. Progression to septic shock involves persistent hypotension and multi-organ failure, with mortality rates above 40% [2]. Globally, sepsis affects approximately 49 million people annually, causing 11 million deaths, representing 20% of worldwide mortality [3]. Low- and middle-income countries (LMICs) bear 85% of sepsis-related deaths due to limited diagnostic and therapeutic resources [4]. In high-income countries, sepsis management incurs significant costs, with the U.S. spending approximately $24 billion annually [5]. Timely diagnosis and antimicrobial therapy are critical, as each hour’s delay increases the mortality risk by 7.6% [6]. Sepsis is predominantly caused by aerobic bacteria, which significantly contributes to its high morbidity and mortality. *Staphylococcus aureus* accounts for 20–30% of BSIs, often linked to invasive procedures like catheter use or endocarditis, with methicillin-resistant strains (MRSA) complicating treatment [7]. *Escherichia coli*, responsible for 25–30% of BSIs, typically originates from urinary or intra-abdominal infections, with increasing resistance to third-generation cephalosporins [8]. *Klebsiella pneumoniae* (10–15% of BSIs) is associated with hospital-acquired pneumonia and exhibits growing carbapenem resistance, with mortality rates for resistant strains reaching 40–50% [9]. *Pseudomonas aeruginosa, Acinetobacter baumannii*, and *Enterococcus* spp. Others are less common (approximately 10% of cases) and pose challenges due to their intrinsic resistance to multiple antibiotics, including carbapenems, making it difficult to treat critically ill patients [10,11]. These aerobic pathogens, particularly Gram-negative bacteria, thrive in the iron-rich environment of blood, evading host defenses through mechanisms like biofilm formation [12]. The rise of antimicrobial resistance (AMR) in these organisms exacerbates treatment challenges, necessitating the development of rapid diagnostics and targeted therapies to improve outcomes [7].

Traditional blood culture remains the cornerstone for diagnosing BSIs, involving the incubation of patient blood samples in nutrient-rich media to detect microbial growth [13]. Automated systems, such as BD BACTEC, BacT/ALERT, and VersaTREK, enhance detection by monitoring CO_2_ production or pH changes, flagging positive cultures within 24–72 hours [14]. Positive samples undergo subculturing, Gram staining, and further identification via biochemical tests or MALDI-TOF, with antimicrobial susceptibility testing (AST) requiring an additional 24 hours[15]. Despite their reliability, these systems face significant challenges. They are time-consuming, often taking 2–5 days for complete pathogen identification and susceptibility profiling, delaying targeted therapy [15]. High costs of commercial media, compounded by supply chain issues and stockouts, limit accessibility, particularly in LMICs [16,17]. Contaminated cultures lead to false positives, resulting in increased hospitalization costs of $6,715–$111,627 per patient [18]. Additionally, proprietary formulations lack customization for advanced molecular diagnostics, such as mNGS, hindering protocol-specific adjustments [19]. The limitations of commercial blood culture media, including high costs, limited availability, and a lack of customization, necessitate the development of reformulated, low-cost, and adaptable alternatives. In low- and middle-income countries (LMICs), the high cost of imported bacterial blood culture commercial media restricts access, particularly in rural areas, due to constrained budgets and supply chain challenges [16,19]. Commercial media stockouts, exemplified by the 2024 BD BACTEC™ shortage, hinder timely sepsis diagnosis [17]. Additionally, proprietary formulations lack flexibility for protocol-specific adjustments required for advanced diagnostics, such as next-generation sequencing (NGS), as additives like resins alter nutrient dynamics and complicate molecular applications [13,20]. In-house, customizable media can be tailored to enhance pathogen detection, optimize DNA retrieval, and neutralize inhibitors, improving diagnostic sensitivity [21]. Affordable formulations promote antimicrobial stewardship, address antimicrobial resistance (AMR), and expand access in resource-limited settings, ultimately enhancing sepsis management and patient outcomes [16]. However, the central question arises, “*Which commercial media can be reformulated to be compatible for culturing bloodstream pathogens?”*. In this study, BHI medium has been selected for re-formulation purposes. *Why*, because BHI provides essential nutrients like amino acids, peptides, and carbohydrates, creating an optimal environment for both fastidious and non-fastidious bacteria [22]. Unlike other common bacterial culture media, such as Luria-Bertani (LB), Tryptic Soy Agar (TSA), Columbia Blood Agar, or any other selective media, BHI’s broad-spectrum support makes it ideal for routine blood culture systems, accommodating diverse microbial metabolisms. Compared to BHI’s enriched composition, which better mimics the blood’s nutrient profile, enhancing pathogen recovery in complex samples [22–24]. Its cost-effectiveness is particularly advantageous in LMICs, where commercial media are prohibitively expensive, enabling its demands for culturing bloodstream pathogens [22]. Additionally, BHI’s flexibility allows for customization of advanced diagnostics, such as mNGS, by optimizing nutrient dynamics and neutralizing inhibitors, thereby addressing the limitations of proprietary commercial media [20]. Thus, BHI’s universal applicability and affordability raise the question of its reformulation compatibility with automated systems, such as BD BACTEC™, facilitating rapid growth detection and feasibility for downstream molecular diagnostics, including qPCR and NGS. Therefore, in this study, we aim to investigate whether in-house reformulation of BHI medium can support effective growth of common aerobic pathogens involved in bloodstream infections directly from blood, and secondly, whether the medium is applicable to molecular diagnostics approaches.

## Materials and Methods

### Ethics statement

In this study, human blood samples were utilized to simulate hospital-grade blood cultures typically obtained from patients. These samples were voluntarily donated by healthy adult individuals who provided informed consent prior to collection at the Department of Biotechnology, University of Inland Norway. The study does not intend to use or analyze human DNA; therefore, any sequencing data originating from the human genome were discarded.

### Bacterial strains

In this study, the bacterial strains selected are common aerobic species known to cause bloodstream infections, such as *Escherichia coli* NCTC 13441, *Staphylococcus aureus* CCUG 17621, *Klebsiella pneumoniae* CCUG 225T, *Acinetobacter baumannii* CCUG 19096T, *Pseudomonas aeruginosa* CCUG 17619, and *Enterococcus faecalis* CCUG 9997. All bacterial strains were revived from glycerol stock and cultivated in BHI agar medium at 37°C for 24 hours prior to isolation of single colony isolates.

### BHI-Blood+: Reformulation of BHI and broth preparation

To enhance the growth of bacteria in Blood, Brain Heart Infusion Broth (BHI) media (53286, Sigma-Aldrich) was supplemented with yeast extract (Y1625, Sigma-Aldrich), L- (−)-Norepinephrine (+)-bitartrate salt monohydrate (norepinephrine) (A9512, Sigma-Aldrich) vitamin mix (MBD0063, Sigma-Aldrich) and polyanetholesulfonic acid sodium salt (SPS) (P2008, Sigma-Aldrich). In-house reformulated BHI was prepared in a two-step process: (1) BHI powder was dissolved in distilled water according to the manufacturer’s instructions. Prior to autoclaving, yeast extract was added at a final concentration of 0.5%. After the autoclave, the media were acclimatized to room temperature. Norepinephrine and SPS stock were prepared according to the manufacturer’s instructions. Norepinephrine and SPS were sterile-filtered through a 0.1 μm syringe filter to prevent mycoplasma contamination. Finally, norepinephrine (final concentration, 300 µM), vitamin mix (final concentration 2%, comprises Biotin : 2 mg/L, Folic acid:2 mg/L, Pyridoxamine-HCl: 10 mg/L, Thiamine-HCl x 2 H2O: 5mg/L, Riboflavin: 5mg/L, Nicotinic acid: 5mg/L, D-Ca-pantothenate: 5mg/L, Cyanocobalamine:0.1mg/L, p-Aminobenzoic acid: 5mg/L, Lipoic acid:5mg/L, KH_2_PO_4_:900mg/L), and SPS (final concentration, 0.05%) were added to the BHI-Yeast broth. For the scope of this literature, this reformulated, in-house developed BHI media is termed BHI-Blood+.

### Bacterial planktonic growth kinetics in BHI-Blood+

Bacterial growth kinetics were measured in BHI Blood+ compared to standard BHI medium to evaluate the planktonic growth performance of various bacterial strains. Bacterial growth kinetics parameters, including lag time and growth rate, were measured as corresponding parameters using the standard microdilution method. Before growth kinetics measurements, bacterial strains were prepared accordingly to the following steps. Bacterial strains were revived from glycerol stock and isolated as single colonies from an overnight static growth culture on BHI agar. Individual colony from each strain was inoculated in 3 mL of BHI broth and cultivated for 2-3 hours with moderate shaking at 37°C, depending on the strain’s background. After brief cultivation, the culture samples were centrifuged at 5000 rpm for 10 minutes. The supernatant was discarded, and the pellets were resuspended in PBS. Bacterial cell density was adjusted to 10^5^ CFU/ml and seeded into each well of a 96-well plate at the final concentration. The Breath-Easy sealing membrane (Z380059, Merck) was used to seal the plates, which were then incubated at 37°C in a Synergy H1 microplate reader. Absorbance at 600 nm for each well was measured at an interval of 30 min over 24 hours. Acquired growth data was analyzed using QurvE software v1.1 [25]. Raw data generated from the microplate reader were transformed to fit into QurvE, and parameters such as growth OD threshold, time at 0 hours (t_0_), and maximum time for growth (t_max_) were set during a quality check on the raw growth curve. The growth threshold was set to 1.5, and t_max_ to 24 hours to extract growth kinetics parameters. Growth profiling of individual bacteria was performed using a growth-fitting exponential growth model with a heuristic linear regression method on log-transformed data [26]. This model determines maximum growth rates (µ_max)_ for individual bacteria cultivated in growth medium. A linear fit was performed using an R2 threshold of 0.95, a Relative standard deviation (RSD) of 0.1, and a dY threshold of 0.05 to estimate the difference in maximum growth and minimum growth. Maximum growth rate (µ_max_) and lag time (λ) were extracted from the linear growth profiling model. µ_max_ was validated using a Gompertz parametric fit [27], and lag time λ was validated using a spline fit model [27].

### Profiling bacterial static growth in BHI-Blood+

Blood samples were collected from healthy volunteers via the sterile venipuncture method. Samples were immediately prepared for culturing to avoid any contamination. Two parallel experimental conditions were established to evaluate bacterial growth dynamics: one using the in-house formulated blood culture medium BHI-Blood+ and the other employing commercially sourced BD BACTEC™ Plus Aerobic/F culture vials (Becton Dickinson and Company, Franklin Lakes, NJ, USA). Time-Resolved Droplet-based CFU Quantification method was introduced to profile the static growth performance of all 6 strains and compare with a commercial standard blood culture medium, BD BACTEC™ PLUS Aerobic/F culture bottles. Before spiking blood, all reference strains were revived from glycerol stock, and an individual isolated colony was inoculated in 3 mL of BHI. Inoculated strains were incubated at 37°C for 2-3 hours with mild agitation to achieve the early exponential growth phase. After short cultivation, culture samples were centrifuged at 5000 rpm for 10 min. Supernatant was discarded to remove standard BHI medium contaminants and any extracellular growth factors. Pellets were washed and resuspended in PBS. The optical density (OD) of each bacterial culture was quantified at 600 nm using a UV-3100PC Spectrophotometer (VWR Life Science, USA), and the CFU/ml was adjusted to a concentration that resulted in a final concentration of 10^2^ CFU/ml when spiked into the blood. The spiked bacterial concentration was verified with the serial dilution colony-forming unit (CFU) method before spiking. 5 mL of donated human blood, spiked with bacteria, was added to 25 mL of the formulated medium and BD BACTEC™ PLUS Aerobic/F culture bottles under sterile conditions. Culture bottles were incubated at 37°C with mild agitation to ensure sufficient oxygen exposure and homogeneity. The incubation time varied based on bacterial type: for *E. coli, K. pneumoniae, A. baumannii*, and *E. faecalis*, samples were collected at 2 and 4 hours, whereas *P. aeruginosa* and *S. aureus* required sampling at 2, 4, and 6 hours due to their comparatively slow growth phenotypes within these experimental parameters. The early time points were chosen to assess growth performance between the commercial BACTEC™ PLUS Aerobic/F culture and BHI-Blood+ and the ability to track early cell proliferation. At each targeted time point, 2 mL samples were aseptically taken from each culture bottle. These samples underwent serial dilution in sterile PBS. 10µl droplets with replicates from each dilution were plated on standard BHI-agar plates and incubated overnight at 37°C. The colonies were later counted to calculate the CFU/mL at each time point, allowing for a quantitative evaluation of the time-resolved bacterial growth progression. Both short-incubation and cultured blood samples were immediately stored at -20°C for use in the downstream experimental method.

### Molecular detection of pathogen in BHI-Blood+

#### DNA extraction

Blood samples spiked with bacterial strains were collected from BHI-Blood+ at specific incubation points: (a) 4 hours for *E. coli, K. pneumoniae, A. baumannii*, and *E. faecalis*, and (b) 6 hours for *P. aeruginosa* and *S. aureus*. Additionally, pure isolates and 24-hour cultures were also subjected to DNA extraction and these samples were used as qualitative controls for the early detection strategy. From each of the 2 ml collected samples, DNA was extracted using the QIAamp Biostic Bacteremia DNA Kit (QIAGEN, Hilden, Germany) according to the manufacturer’s instructions. DNA was eluted in nuclease-free water and quantified with the Qubit dsDNA HS Assay Kit on a Qubit 4 Fluorometer (Thermo Fisher). The purity of DNA samples was checked using the NanoDrop ND-1000 Spectrophotometer (Thermo Fisher) to measure the OD 260/280 ratio of 1.8 and the OD 260/230 ratio of 2.0–2.2. If necessary, optional concentration and purification were performed with the Agencourt AMPure XP system (Beckman Coulter, USA). Extracted DNA samples were subjected to qPCR-based pathogen detection, followed by Oxford Nanopore.

#### Real-time polymerase chain reaction

To detect bacteria in the extracted DNA samples, Real-time polymerase chain reaction (qPCR) was utilized. In this study, species-specific primers were used to detect species that were spiked in blood. Detailed primer sequences and references are listed in the supplementary Table S1. qPCR were performed on a 7500 Fast Real-Time PCR System (Applied Biosystems, Thermo Fisher Scientific, Waltham, MA, USA), employing the following thermal profile: an initial denaturation stage at 95°C for 12 minutes to activate the Hot FIREPol enzyme, followed by 40 cycles comprising denaturation at 95°C for 25 seconds, annealing at 60°C for 45 seconds, and an extension phase at 72°C for 1 minute. Cycle threshold (Ct) values were recorded for each reaction and used for qualitative evaluation of the presence of bacterial DNA in extracted samples. Mean Ct values calculated from replicates and visualized differences using an in-house developed R shiny app. The interpretation of early detection was performed based on delta CT differences between 4/6 hours samples, pure isolated, and 24-hours culture.

#### Oxford nanopore sequencing

In this study, Oxford nanopore sequencing was conducted on 4 or 6-hour culture samples to evaluate possibility of early detection of spiked species and whether BHI-Blood+ can be used as a routine blood culture medium for detecting bacteria using a next-generation sequencing platform AMPure XP-purified DNA samples were subjected to library preparation using the Rapid Barcoding Kit 96 V14 (SQK-RBK114.96, Oxford Nanopore Technologies, Oxford, UK), which facilitated the multiplexing of samples by incorporating distinct barcodes into each DNA sample. The library preparation was carried out according to the manufacturer’s protocol without any modifications. The prepared sample library was loaded onto the MinION Flow Cell (R10.4.1, FLO-MIN114, Oxford Nanopore Technologies, Oxford, UK) and sequenced using the MinION MK1D device. The parameters for the sequencing run were set to Fast model v4.3.0 basecalling, implemented by Dorado 7.6.8, with a Minimum Q score of 7 and a run time of 24 hours. The output files generated by the MinKNOW software were analysed using minimap2 v2.28-r1209 based alignment to the reference genome. The number of reads and genome coverage of spiked strains were estimated to confirm the detection and identity of the spiked bacterial species. The reference assemblies for the strains used in this study can be found in the NCBI database by using the accession numbers *E. coli* NCTC 13441 (GCA_900448475.1), *P. aeruginosa* CCUG 17619 (GCA_024507955.1), *K. pneumoniae* CCUG 225T (GCA_000742135.1), *A. baumannii* CCUG 19096T (GCA_900444725.1), *S. aureus* CCUG 17621(GCA_028596245.1), *E. faecalis* CCUG 9997 (GCA_022406475.1). Reads aligned to reference bacterial genomes were subjected to iterative BLASTN searches to confirm the species and identify false positives.

### Statistical analysis

Statistical analysis on both growth profile parameters was performed using GraphPad Prism (version 10.5.0). A non-parametric Mann-Whitney U test was applied to both planktonic and static bacterial growth profiles. A two-tailed p value of <0.01 was considered statistically significant.

## Results

### Impact of BHI-Blood+ cocktail on bacterial planktonic growth: No detrimental effect observed

Figure 1 (A) illustrates an overview of the experimental procedure to perform the planktonic growth profile of all 6 strains in BHI Blood+. Profiling *E. coli* planktonic growth under various conditions provides a deeper understanding of the effects of norepinephrine, a vitamin mix, and an SPS cocktail. Growth curves, depicted as log growth (OD600) against time, reveal distinct phases: an initial lag phase during which bacterial adaptation occurs, followed by an exponential growth phase that peaks approximately 2-8 hours later, and a subsequent stationary phase beginning at 9 hours and continuing thereafter Figure 1 (B). When the two conditions, BHI and BHI-Blood+, are compared, the curves of both conditions show overlapping trajectories, indicating no significant deviation in growth patterns. Quantitative analysis of the lag phase and the maximum growth rate (µ_max_), with statistical significance (p < 0.01) between BHI and BHI-Blood+ conditions, shows no significant differences between conditions and therefore supports the absence of inhibitory effects exerted by BHI-Blood+ additives Figure 1 (C).

**Figure 1.**
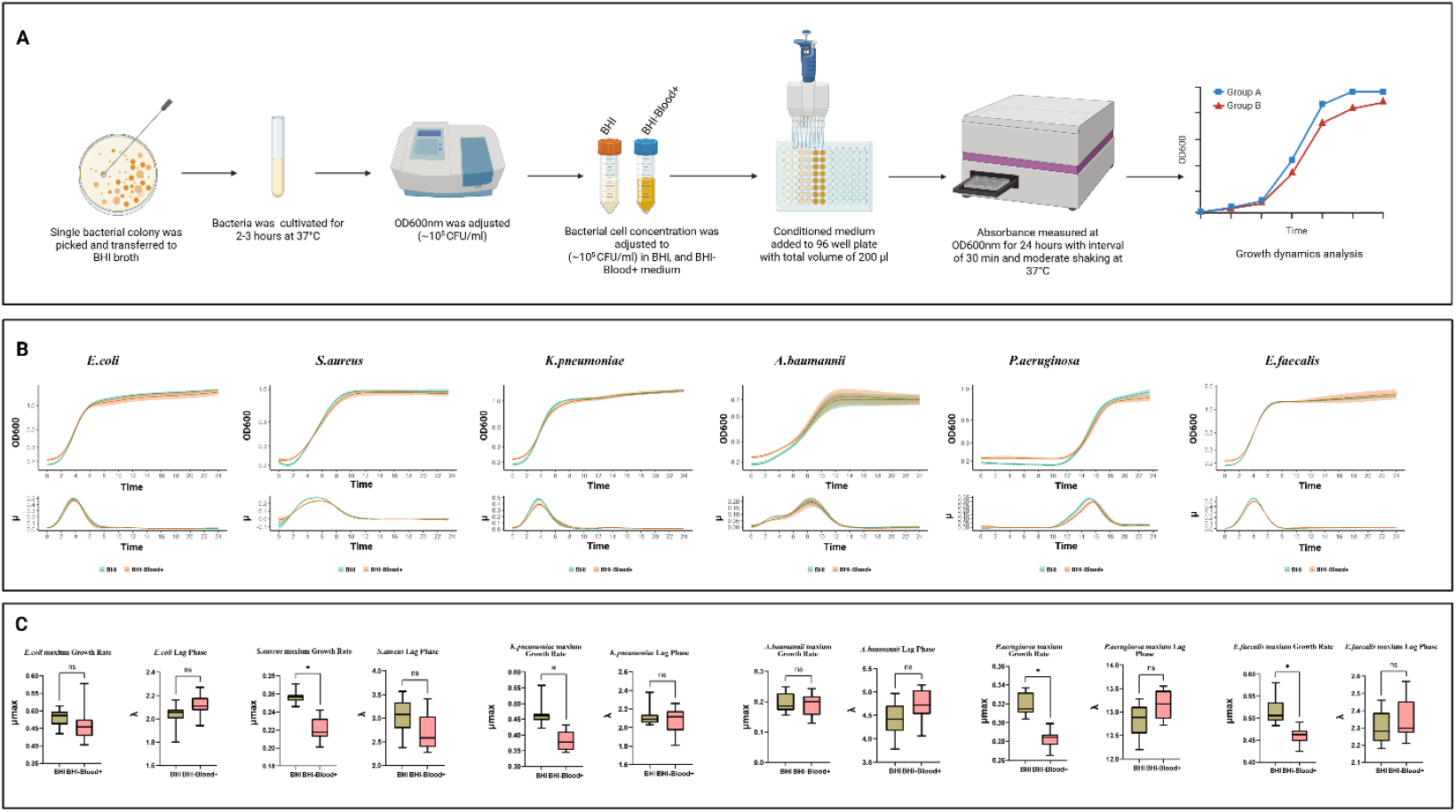
An in-depth analysis of bacterial growth dynamics of six species, including *Escherichia coli, Staphylococcus aureus, Klebsiella pneumoniae, Acinetobacter baumannii, Pseudomonas aeruginosa*, and *Enterococcus faecalis*, under two culture conditions: (A) Overview of experimental workflow for bacterial planktonic growth dynamics in BHI and BHI-Blood+ supplemented with norepinephrine, vitamin mix, and SPS. (B) Each panel includes a growth curve representing planktonic growth over 24 hours, (C) a growth rate distribution, and a lag phase, as analyzed by QurvE on log-transformed data. Box plots indicate statistical comparisons with no significant (ns) labels, where p ≤ 0.01 and significant differences are marked with asterisks (*)

*S. aureus*, on the other hand, exhibits a distinct growth dynamic compared to *E. coli* when reacting to the additives of BHI-Blood+ Figure 1 (C). While the lag phase shows no significant differences between BHI and BHI-Blood+, the maximum growth rate (µ_max_) of *S. aureus* under BHI conditions dominates that of BHI-Blood+. Growth rate results also support these findings, indicating that a lower shift in µ_max_ occurred after 5 hours. While the growth rate of BHI peaked at late 4 hours, both conditions overlapped in the exponential phase between 2 and 8 hours, Figure 1 (B). These findings suggest that while the additives do not alter the lag phase, the reduction in µ_max_ in BHI-Blood+ may be due to a metabolic adaptation in the presence of these additives.

A similar trend of growth trajectories was also observed for *K. pneumoniae*, Figure 1 (B). Similar adaptation periods were observed across both conditions, supporting non-significant differences in the lag phase, as shown in Figure 1 (C). However, the µ_max_ rates showed a significant reduction in BHI-Blood+ compared to BHI, suggesting that the peak growth capacity of *K. pneumoniae* is impaired by the additives, Figure 1 (C). Additionally, the growth curves for *K. pneumoniae* display an exponential growth phase that overlapped in both conditions, supporting a minimal effect of additives on bacterial growth kinetics Figure 1 (B). *E. Faecalis* demonstrates similar trends to *S. aureus* and *K. pneumoniae* in showing differences in µ_max,_ Figure 1 (C). However, there were no significant differences in the lag phase. In contrast, the growth behavior of *A. baumannii* exhibited no significant difference in either the lag phase or growth rate, implying that the additives do not alter its peak planktonic growth capacity, Figure 1 (C). These findings are also supported by a biphasic exponential growth pattern, overlapping between both conditions in *A. baumannii* (exponential phases, 2-4 and 4-6 hours), which depicts consistent metabolic adaptation across conditions, Figure 1 (B). On the other hand, compared to all other strains, P. aeruginosa demonstrates a prolonged lag time when cultivated with both media, Figure 1 (B), while significantly differing from each other in maximum growth rate, Figure 1 (C).

### Compatibility between BHI Blood+ medium and BD BACTEC for short cultivation

Figure 2 (A) depicts Bacterial Time-Resolved Droplet-based CFU Quantification assessment of static growth profile of six clinically significant pathogens: *E. coli, K. pneumoniae, P. aeruginosa, E. faecalis, A. baumannii*, and *S. aureus*, in two culture media: BHI-Blood+ and BACTEC™ PLUS Aerobic/F culture. CFU quantification results suggest that *E. coli*, with average values of ∼10^5^ CFU/mL at 2 hours and 10^6^ CFU/mL at 4 hours, highlights the in-house medium’s equivalent performance to BD BACTEC™, without any significant difference between the two media Figure 2 (B). Similarly, *K. pneumoniae* and *E. faecalis* grew rapidly over a 4-hour period Figure 2 (B). Starting at 10^2^ CFU/mL, the BHI-Blood+ supported growth to 10^5^ CFU/mL by 2 hours and 10^6^ CFU/mL by 4 hours, while BACTEC™ PLUS Aerobic/F culture media achieved similar CFU/mL compared to BHI-Blood+ without statistical significance, confirming the efficacy of BHI-Blood+ for the early detection of these pathogens. However, *A. baumannii* demonstrated ∼10^3^ CFU/mL at 2 hours and increased to ∼10^5^ CFU/mL by 4 hours Figure 2 (B), which was slower static growth as compared to *E. coli, K. pneumoniae, and E. faecalis*, but when compared to BACTEC™ PLUS Aerobic/F culture media, BHI-Blood+ also showed its compatibility for *A. baumannii* growth in blood and no significant differences.

**Figure 2:**
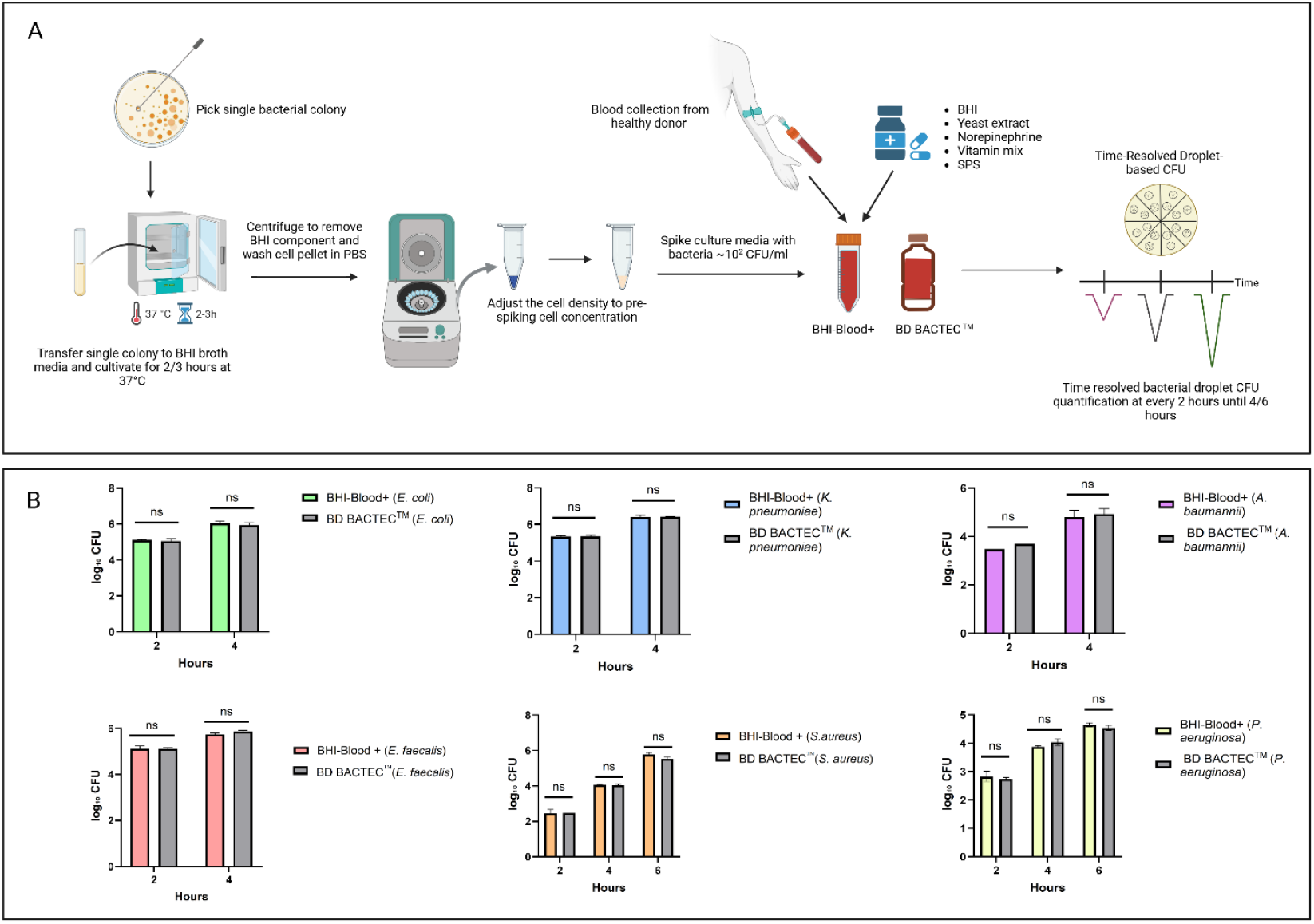
(A) illustrates experimental overview of profiling bacterial static growth dynamics. (B) The growth dynamics of 4 bacterial species – *E. coli, K. pneumoniae, E. faecalis, A. baumannii*– were evaluated using BHI-Blood+ and BD BACTEC™ PLUS Aerobic/F culture media over a period of 2 and 4 hours, whereas *P. aeruginosa and S. aureus* for 2,4, and 6 hours, with log_10_ CFU/ml counts measured at intervals of 2 hours. Statistical analysis showed no significant difference (ns) between the two conditions at each time.

On the other hand, the delayed static growth dynamics of both *P. aeruginosa* and *S. aureus* were assessed at an additional time point, 6 hours Figure 2 (B). *P. aeruginosa* growth progressed over 6 hours, starting at ∼10^3^ CFU/mL at 2 hours, slowly rising to 104 CFU/mL by 4 hours, and reaching ∼10^5^ CFU/mL by 6 hours in both media. Contrary, *S. aureus* showed limited proliferation at 2 hours (∼10^2^ CFU/mL) in both media, increasing to the 104 CFU/mL range by 4 hours. By 6 hours, the BHI-Blood+ supported a higher bacterial load (∼10^6^ CFU/mL), with no statistical differences compared to BACTEC™ PLUS Aerobic/F culture media Figure 2 (B).

These results demonstrate that BHI-Blood+ and BD BACTEC™ PLUS Aerobic/F culture media support comparable early-phase bacterial proliferation *in vitro* across a wide range of Gram-negative and Gram-positive organisms, indicating that BHI-Blood+ is compatible with bloodborne pathogens.

### BHI Blood+ supports bacterial early detection

Figure 3 (A) demonstrates molecular detection strategy for pathogen spiked in both BHI-Blood+ and BD BACTEC™ PLUS Aerobic/F culture media. DNA extracted from blood cultures spiked with six sepsis-relevant bacterial strains (*E. coli, P. aeruginosa, K. pneumoniae, A. baumannii, S. aureus*, and *E. faecalis*) using the BiOstic protocol showed high purity via NanoDrop measurements, confirming an efficient extraction process. To assess whether BHI-Blood+ supports early detection through short incubation periods, each strain was analyzed at specific time points: *E. coli* (4 and 24 hours), *P. aeruginosa* (6 and 24 hours), *K. pneumoniae* (4 and 24 hours), *A. baumannii* (4 and 24 hours), *S. aureus* (6 and 24 hours), and *E. faecalis* (4 and 24 hours). Mean Ct value differences were evaluated to infer detection of bacterial proliferation in spiked samples.

**Figure 3:**
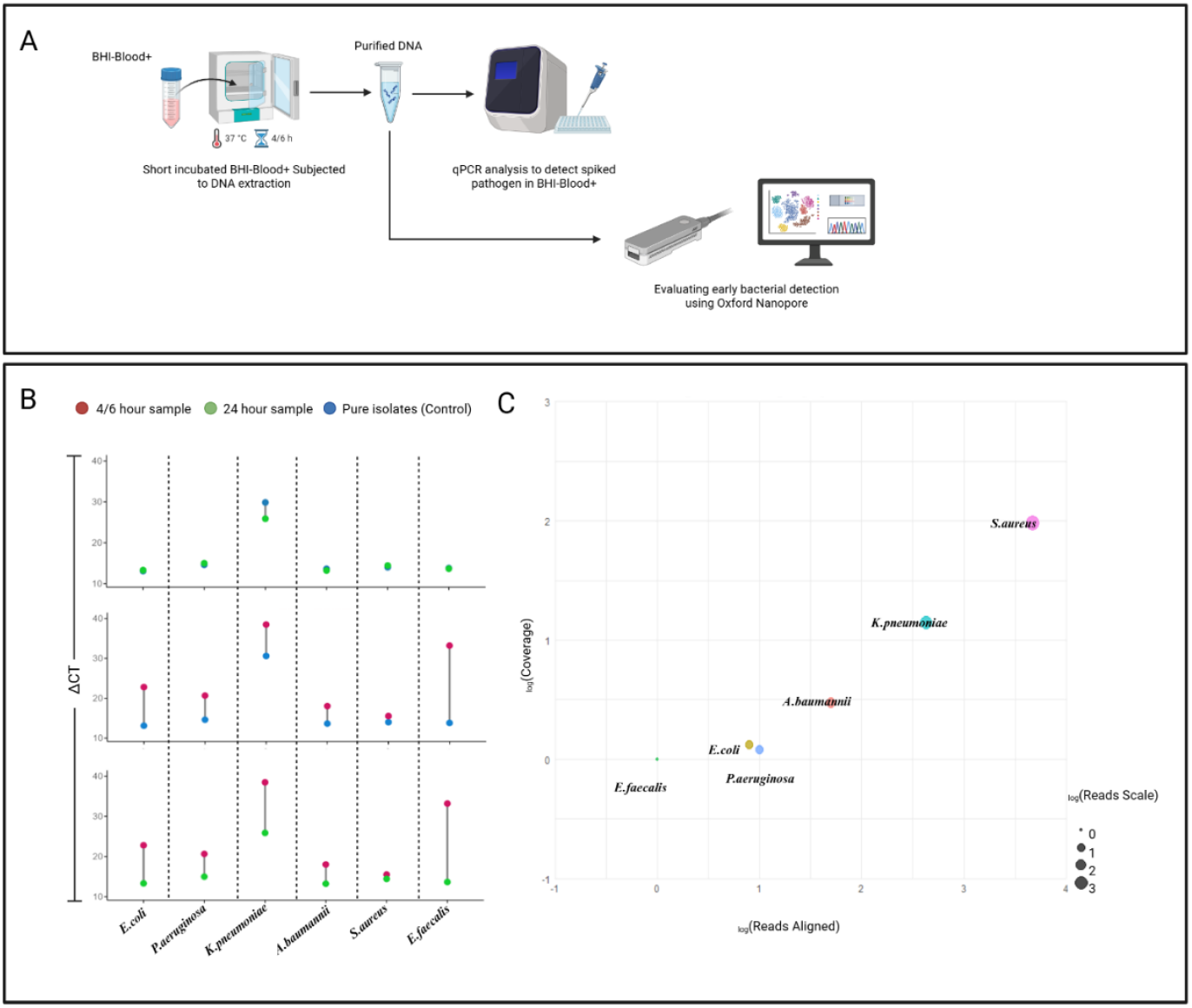
(A) Illustrates experimental workflow for molecular detection of targeted species in BHI-Blood+ medium. (B) Depicts the analysis of species-specific detection across different samples spiked by bacterial strains. ΔCt differences between control and culture conditions demonstrate earlier detection efficiency. Samples collected at 4 hours (Red dots), 24 hours (Green dots), and pure isolates (Blue dots). (C) displays a bubble plot illustrating the relationship between log-transformed sequencing reads aligned to reference genome (X axis = _log_Reads) and Coverage (Y= axis _log_Coverage). The bubble size (based _log_Reads scale) represents the correlation between read count and coverage, with larger bubbles indicating a higher ratio of read count to coverage, signifying enhanced species detection.

Results, summarized as Ct values in Table S2-6, indicate that all six strains were detected at their respective time points after short incubation. Delta Ct values between short incubation and positive controls showed early detection for all strains except *E. faecalis*, which exhibited higher delta Ct values. Detection template generated from pure isolates, and 24-hour sample defines the early detection efficiency. As 24-hour samples provided sufficient biomass for detection of all species, reflected by the lowest delta Ct values when compared to pure isolates. Comparison of short incubation and 24-hour cultures revealed trajectories sufficient to confirm cellular proliferation of these strains earlier than 24 hours.

On the other hand, minimap2 analysis reports the number of reads aligned to reference genome derived from nanopore sequencing and significant *E-*value to designated species validates species detection (summarized in Table S7). Sequencing results confirm the detection of all targated species except *E. faecalis*, and the reason for this detection failure is still unclear. However, results from both planktonic and static growth profile suggest that shows that errors may arise during DNA extraction or sample library preparation, which would be difficult to trace because the samples were not subjected to host depletion. Among the species, *S. aureus* and *K. pneumoniae* depicts the highest read counts and coverage. In contrast, lower read counts for *E. coli, P. aeruginosa*, and *A. baumannii* indicate potential technical issues, such as low quality of sequencing due to short-reads generated by the DNA extraction process itself, higher level of host DNA contamination, and importantly, the DNA extraction process is not optimal for short incubation sample condition, which represents high host DNA compared to low bacterial DNA concentration. Overall, both qPCR and nanopore sequencing validate the compatibility of BHI-Blood+ media for early bacterial detection.

## Discussion

### BHI-Blood+: A cost-effective alternative for blood culture diagnostics in resource-limited settings

Blood is generally considered a sterile environment, free from viable pathogens, under healthy conditions. However, commensal microbes can be translocated transiently and sporadically from different body sites into the bloodstream [28]. Even a few percent of pathogens that reach the bloodstream, opsonization and phagocytosis-based immune responses actively suppress the growth of these pathogens. These host defense mechanism releases multiple inhibitors to create a hostile environment for pathogens [29]. Commercial blood culture media are formulated in such a way that they not only neutralize the inhibitors released by host immune responses but also provide nutrients and growth factors to support the recovery of pathogens, facilitating accurate sepsis diagnosis. This culture system is regarded as the gold standard for detecting bloodstream infections. BD BACTEC™ comprises ingredients, which include Soybean-Casein digest broth, yeast extract, amino acids, sugar, vitamins, SPS, antioxidants/reductants, nonionic adsorbing resin, and cationic exchange resin, which effectively support the growth of sepsis-inducing pathogens. While these systems support the growth of pathogens, they are yet limited by high costs and supply shortages [30]. For instance, the cost of BD BACTEC™ Plus Aerobic/F bottles typically ranges from $10– 15 per bottle (30 ml), depending on the suppliers and import expenses in low-resource environments [31]. Addressing these limitations of commercial blood culture media, the necessity of an alternative has opened a door to reformulating the existing BHI media. This study evidently demonstrated that use of blood-supported, in-house formulated brain heart infusion (BHI-Blood+) offers similar growth dynamics efficacy of sepsis inducing aerobic pathogens, thus receiving attention as a potential alternative to commercial media. One big advantage of BHI-Blood+ is that, besides utilizing locally sourced BHI, yeast extract, and other additives, it is estimated at $4–5 per bottle (30 ml). Such a significant cost reduction enhances accessibility in resource-constrained settings without compromising bacterial growth performance. Additionally, the additives in BHI-Blood+ comprise 4-5 ingredients (yeast extract, norepinephrine, vitamin mix, and SPS), compared to 8-9 ingredients in BD BACTEC™ Plus Aerobic/F bottles, offering an attractive option for hospitals and clinics in developing countries, where budget constraints often limit access to advanced diagnostic tools.

On the other hand, the individual supplements added in BHI-Blood+ are classical requirements for bacterial growth. Starting with yeast extract, which provides essential amino acids, peptides, and vitamins, supporting the metabolic needs of individual bacteria [32]. Vitamin mix used in BHI-Blood+ is to supplement additional cofactors to promote bacterial growth in BHI-Blood+ [33–35]. However, one of the most essential addon in BHI-Blood+ is the norepinephrine, a group of hormones and neurotransmitters known as Catecholamines. Catecholamines are released into the bloodstream in response to stress, regulating multiple physiological functions, particularly by escalating blood pressure and increasing cardiac output [36,37]. Notably, some studies have depicted the release of catecholamines in the bloodstream as the body’s acute response to infection and play an important role in maintaining vascular tone, heart rate, and blood pressure to counteract the hypotension and tissue hypoperfusion characteristics during septic shock [36]. However, some studies have also shown that catecholamines, especially norepinephrine, are found to promote bacterial growth by facilitating the acquisition of iron from the host through interaction with host proteins such as transferrin. For example, it has been reported that norepinephrine promotes the proliferation and virulence of pathogenic bacteria in the iron-limited environment [38–41]. Besides, blood can also be considered an iron-limiting environment, particularly during infection or inflammation. Hepcidin, a hormone, plays a critical role in reducing the amount of iron entering the bloodstream during infection and inflammation, which leads to starvation of pathogens of iron in the bloodstream [12,42,43]. Therefore, during bacterial culture in blood, catecholamines, especially norepinephrine, were supplemented in BHI-Blood+ to accelerate bacterial growth. Moreover, SPS is a cornerstone component in BHI-Blood+, enhancing pathogen recovery by neutralizing host inhibitors and preventing clotting[44]. Thus, the synergistic effect of all these components in BHI-Blood+ creates an optimal growth environment for bloodborne bacteria. This makes it a viable option for clinical microbiological laboratories, especially in resource-limited environments. Due to its open formulation, it has the potential to further customize BHI-Blood+, thereby enhancing its demand. Unlike standardized and exclusive commercial media, BHI-Blood+ can be adapted to specific local needs. For example, laboratories can adjust supplements to optimize the growth of specific pathogens present in their region [45]. Additionally, BHI-Blood+ offers an advantage in terms of ease of preparation, accessibility, and feasibility in the supply chain. BHI-Blood+ can be easily prepared in laboratories using standard equipment and locally available materials, such as individual components of BHI or commercially available, pre-prepared BHI powders, and additives like vitamin mix, norepinephrine, and SPS. This local production minimizes dependence on international suppliers, which often face delays or disturbances in low-resource environments due to logistical challenges or import restrictions. With such consistent local production, ensures a steady supply and flexible volume preparation tailored to the needs of internal diagnostics or research and thereby supporting uninterrupted diagnostic services. Moreover, unlike BD BACTEC™ or any commercial automated systems which demand specialized equipment and skilled staff, preparation of BHI-Blood+ does not require significant investments in technology or training. Its adaptability enables labs to scale production to meet diverse demands, independent of pre-packaged commercial products. [46].

### BHI-Blood+ is compatible with rapid sepsis diagnostics and reproducible host-pathogen research

Traditional blood culture strategies require 24-72 hours for detectable bacterial growth, often making them too slow for critical cases. Thus, delayed treatment in sepsis can cause an increase in mortality risk of 7-8% per hour. A study done by Jawad et al demonstrated that shorter blood culture can be achieved with BD BACTEC™ and thus fast forward blood sepsis diagnosis from sample collection to identify pathogens and ARGs within a 7–9-hour timeframe [47]. This study has opened a new dimension of possibilities, revealing that short cultivation can be a novel strategy for diagnosing blood sepsis and enhancing the early detection of pathogens in the history of blood sepsis. As the BD BACTEC™ system supports a short cultivation strategy, BHI-Blood+ supplemented with norepinephrine, vitamin mix, and SPS sheds light on it as a promising medium for short-term incubation, demonstrated in this study. All pathogens used in this study have been shown to adapt to the short-term cultivation strategy. This study has shed light on the possibility of earlier detection of bacteria through a simple cultivation strategy. In principle, BACTEC™ Plus Aerobic/F culture vials work as microorganisms in the medium proliferate, CO2 levels rise as a result of the metabolism of substrates present in the medium. The vial’s integrated sensors detect increased CO_2_ levels through enhanced fluorescence, which is apparently monitored by BD BACTEC™ system [48]. The BACTEC FX system can flag positive culture for most bloodstream infection pathogens when the CFU reaches > 10^7^-10^8^ CFU/ml[49], which indicates that detecting certain levels of CO_2_ can take more than 12 hours or even longer, even if bacterial growth is in the exponential phase. This study has reproduced the findings of Jawad et al [47] that the early detection of pathogens can be done by tracking their early proliferation in culture medium. As compared to BD BACTEC™, BHI-Blood+ demonstrated its compatibility for early detection at the cellular level at <8 hours. Additionally, the medium’s compatibility with molecular platforms like qPCR and nanopore sequencing positions it as a valuable tool for molecular-based pathogen detection at < 24 hours, and metagenomic studies for pathogen surveillance. Thus, as a low-cost alternative in-house media, BHI-Blood+ demonstrated its potential as a component in rapid sepsis diagnostics research. However, to demonstrate its full potential, challenges remain in evaluating anaerobes and other fastidious pathogens, clinical validation of real patient samples with potential inhibitors (antimicrobials), cost barriers in resource-limited settings, and feasibility in hospital automation and upscaling.

On the other hand, apart from sepsis diagnosis, BHI-Blood+ can also offer another advantage in addressing the necessities of optimizing culture media for reproducible antibiotic sensitivity tests directly in blood cultures [50,51]. Previous studies have demonstrated that blood culture resuscitation in culture media can support direct blood culture susceptibility studies, yielding results comparable to those obtained with standard methods, thereby reducing the time to actionable results [52]. Adding BHI-Blood+ to this extent of capability can be valuable in situations where rapid sensitivity tests are important for patient management. Additionally, antibiotic sensitivity tests using BHI-Blood+ have the potential to support reproducible research in understanding complex host-pathogen interactions in bloodstream infections and their molecular mechanisms of antibiotic resistance. As discussed earlier, BD BACTEC™ is only used for commercial sepsis detection systems, and its use in research is not straightforward, as it is limited to customization or open formulation. The open formulation of BHI-Blood+ provides control over its components for reproducible research. To date, the current understanding of infect-omics of bloodstream pathogens is based on blood agar static cultivation systems. This study has opened new doors for researchers to utilize BHI-Blood+ medium as a routine, low-cost culture system to study the molecular mechanisms of bloodstream pathogens and their interactions with the host. This open formulation can give researchers the advantage of further formulating the medium and upgrading it according to their specific research needs, offering greater flexibility compared to the more restrictive BD BACTEC system.

## Conclusion

In conclusion, BHI-Blood+ addresses key barriers to blood culture diagnostics—cost, accessibility, and customization—while demonstrating its compatibility in the rapid detection of sepsis. Its potential to transform sepsis diagnostic and reproducible research practices into resource-limited settings underscores the need for further validation and optimization to fully realize its capabilities across diverse clinical and research applications.

## Acknowledgements

This study was supported by the OH-AMR-Diag project, funded by the Research Council of Norway (project number 336420).

## Author Contributions

**Resources:** Rafi Ahmad. **Conceptualization:** Rafi Ahmad, Jonathan Hira, and Jawad Ali. **Methodology:** Nasib Bin Mahbub, Jonathan Hira, and Jawad Ali. **Investigation and formal analysis: Jonathan Hira and** Nasib Bin Mahbub and Jonathan Hira. **Supervision:** Rafi Ahmad, Jonathan Hira, and Jawad Ali. **Writing – original draft:** Jonathan Hira. **Writing – review & editing:** Jonathan Hira, Nasib Bin Mahbub, Jawad Ali, and Rafi Ahmad.

