## Supplementary material for "Low-Cost In-House Re-formulated Brain Heart Infusion Medium for Effective Planktonic Growth and Early Detection of Bloodstream Bacterial Pathogens": Table S1

**S1 Table: List of species-specific primers used in this study**

| Target Species | Target | Product size (bp) | Species specific primer | Reference |
|---|---|---|---|---|
| *E. coli* NCTC 13441 | Universal stress protein- *uspA* | ~ 884 | F- 5’ CCGATACGCTGCCAATCAGT 3’<br>R- 5’ ACGCAGACCGTAGGCCAGAT 3’ | [1] |
| *S. aureus* CCUG 17621 | Thermonuclease *-nuc* | ~ 65 | F- 5’ GGGTTGATACGCCAGAAACG 3’<br>R- 5’ TGATGCTTCTTTGCCAAATGG 3’ | [2] |
| *K. pneumoniae* CCUG 225T | Hemolysin *khe* | ~ 486 | F- 5’ TGATTGCATTCGCCACTGG 3’<br>R- 5’ GGTCAACCCAACGATCCTG 3’ | [3] |
| *A. baumannii* CCUG 19096T | DNA gyrase *gyrA* | ~344 | F- 5'-AAATCTGCCCGTGTCGTTGGT -3’<br>R- 5’-GCCATACCTACGGCGATACC -3’ | [4] |
| *P. aeruginosa* CCUG 17619 | Phenazine biosynthesis protein *PhZA2* | ~325 | F- 5’ GTTTACCGACAACCTGGAA 3’<br>R- 5’ GCAATAGCCCTGCGGATAC 3’ | [5] |
| *E. faecalis* CCUG 9997 | heat shock proteins groES | ~185 | F- 5’ GGAATTGTTCTTGCATCCGT 3’<br>R- 5’ ACAATTAAGTATTCTACGCC 3’ | [6] |

**S2: CFU count comparison between BHI-Blood+ and BD BACTEC™ at different time points for all the strains**

| Species | Time (Hour) | Culture medium | Replicate 1 (CFU/ml) | Replicate 2 (CFU/ml) |
|---|---|---|---|---|
| ***E. coli NCTC* 13441** | 2 | BHI-Blood+ | $1.40 \times 10^5$ | $1.20 \times 10^5$ |
| | | BD BACTEC™ | $1.40 \times 10^5$ | $9.00 \times 10^4$ |
| | 4 | BHI-Blood+ | $1.30 \times 10^6$ | $9.00 \times 10^5$ |
| | | BD BACTEC™ | $1.10 \times 10^6$ | $7.00 \times 10^5$ |
| ***K. pneumoniae* CCUG 225T** | 2 | BHI-Blood+ | $2.40 \times 10^5$ | $2.00 \times 10^5$ |
| | | BD BACTEC™ | $2.50 \times 10^5$ | $2.00 \times 10^5$ |
| | 4 | BHI-Blood+ | $3.00 \times 10^6$ | $2.20 \times 10^6$ |
| | | BD BACTEC™ | $2.60 \times 10^6$ | $2.70 \times 10^6$ |
| ***A. baumannii* CCUG 19096T** | 2 | BHI-Blood+ | $3.00 \times 10^3$ | $3.00 \times 10^3$ |
| | | BD BACTEC™ | $5.00 \times 10^3$ | $5.00 \times 10^3$ |
| | 4 | BHI-Blood+ | $4.00 \times 10^4$ | $1.00 \times 10^5$ |
| | | BD BACTEC™ | $1.20 \times 10^5$ | $6.00 \times 10^4$ |
| ***P. aeruginosa* CCUG 17619** | 2 | BHI-Blood+ | $5.00 \times 10^2$ | $9.00 \times 10^2$ |
| | | BD BACTEC™ | $6.00 \times 10^2$ | $5.00 \times 10^2$ |
| | 4 | BHI-Blood+ | $8.00 \times 10^3$ | $7.00 \times 10^3$ |
| | | BD BACTEC™ | $9.00 \times 10^3$ | $1.30 \times 10^4$ |
| | 6 | BHI-Blood+ | $5.00 \times 10^4$ | $4.00 \times 10^4$ |
| | | BD BACTEC™ | $4.00 \times 10^4$ | $3.00 \times 10^4$ |
| ***E. faecalis* CCUG 9997** | 2 | BHI-Blood+ | $1.10 \times 10^5$ | $1.60 \times 10^5$ |
| | | BD BACTEC™ | $1.20 \times 10^5$ | $1.40 \times 10^5$ |
| | 4 | BHI-Blood+ | $5.00 \times 10^5$ | $6.00 \times 10^5$ |
| | | BD BACTEC™ | $8.00 \times 10^5$ | $7.00 \times 10^5$ |
| ***S. aureus* CCUG 17621** | 2 | BHI-Blood+ | $2.00 \times 10^2$ | $4.00 \times 10^2$ |
| | | BD BACTEC™ | $3.00 \times 10^2$ | $3.00 \times 10^2$ |
| | 4 | BHI-Blood+ | $1.10 \times 10^4$ | $1.20 \times 10^4$ |
| | | BD BACTEC™ | $1.20 \times 10^4$ | $1.00 \times 10^4$ |
| | 6 | BHI-Blood+ | $5.00 \times 10^5$ | $7.00 \times 10^5$ |
| | | BD BACTEC™ | $3.00 \times 10^5$ | $4.00 \times 10^5$ |

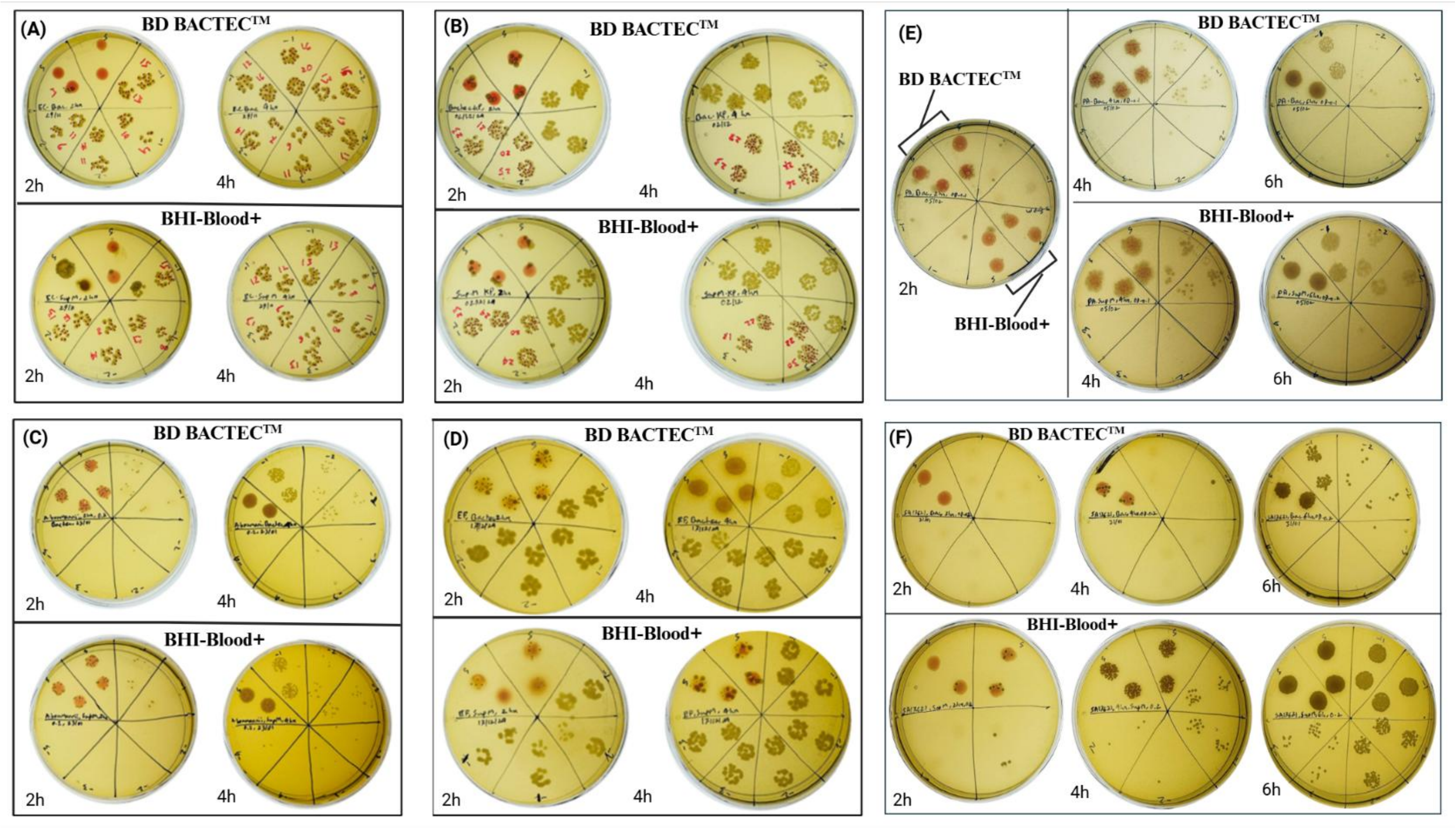


**Figure S1**: Overview of comparative static growth analysis of target species in BHI-Blood+ Media and BD BACTEC™. Colony forming units for individual species are examined on BHI agar plates at different time points (2,4 or 6) after being cultivated at 37°C. (A) *E coli* NCTC13441, (B) *K. pneumoniae* CCUG225T, (C) *A. baumannii* CCUG 19096T, (D) *E. faecalis* CCUG9997 (E) *P. aeruginosa* CCUG17619, (F*) S. aureus* CCUG17621.

**S3 Table: qPCR CT values**

| Species | Timepoint (hours) | R1 | R2 | R3 | Mean Ct | Std. Dev. |
|---|---|---|---|---|---|---|
| *E coli* NCTC13441 | Isolates | 13.07 | 13.09 | 13.1 | 13.09 | 0.02 |
| | 4 | 22.84 | 22.74 | 22.79 | 22.79 | 0.05 |
| | 24 | 13.3 | 13.3 | 13.32 | 13.31 | 0.01 |
| *P. aeruginosa* CCUG17619 | Isolates | 14.6 | 14.6 | 14.6 | 14.60 | 0.00 |
| | 4 | 20.71 | 20.62 | 20.65 | 20.66 | 0.05 |
| | 24 | 14.9 | 15 | 15.06 | 14.99 | 0.08 |
| *K. pneumoniae* CCUG225T | Isolates | 30.9 | 30.3 | NA | 30.60 | 0.42 |
| | 4 | 37.44 | 39.54 | NA | 38.49 | 1.48 |
| | 24 | 25.8 | 26 | 25.85 | 25.88 | 0.10 |
| *A. baumannii* CCUG 19096T | Isolates | 13.63 | 13.62 | 13.64 | 13.63 | 0.01 |
| | 4 | 17.95 | 18.07 | 18.07 | 18.03 | 0.07 |
| | 24 | 13.2 | 13.2 | 13.26 | 13.22 | 0.03 |
| *S. aureus* CCUG17621 | Isolates | 14 | 14 | 14 | 14.00 | 0.00 |
| | 4 | 15.54 | 15.52 | 15.49 | 15.52 | 0.03 |
| | 24 | 14.4 | 14.4 | 14.49 | 14.43 | 0.05 |
| *E. faecalis* CCUG9997 | Isolates | 13.8 | 13.8 | 13.8 | 13.80 | 0.00 |
| | 4 | 34.4 | 32.96 | 32.24 | 33.20 | 1.10 |
| | 24 | 13.6 | 13.7 | 13.65 | 13.65 | 0.05 |

**S4 Table: qPCR ΔCT values between 4 hours and 24 hours**

| Species | Mean_Ct_4 | Mean_Ct_24 | SD_4 | SD_24 | ΔCt |
|---|---|---|---|---|---|
| *A. baumannii* CCUG 19096T | 18.03 | 13.22 | 0.06928203 | 0.03464102 | 4.81 |
| *E. faecalis* CCUG9997 | 33.2 | 13.65 | 1.09981817 | 0.05 | 19.55 |
| *E.coli* NCTC13441 | 22.79 | 13.3066667 | 0.05 | 0.01154701 | 9.48 |
| *K. pneumoniae* CCUG225T | 38.49 | 25.8833333 | 1.48492424 | 0.1040833 | 12.61 |
| *P. aeruginosa* CCUG17619 | 20.66 | 14.9866667 | 0.04582576 | 0.08082904 | 5.67 |
| *S. aureus* CCUG17621 | 15.5166667 | 14.43 | 0.02516611 | 0.05196152 | 1.09 |

**S5 Table: qPCR ΔCT values between pure isolates and 4 hours**

| Species | Mean_Ct_Isolates | Mean_Ct_4 | SD_Isolates | SD_4 | ΔCt |
|---|---|---|---|---|---|
| *A. baumannii* (CCUG 19096T) | 13.63 | 18.03 | 0.01 | 0.06928203 | 4.4 |
| *E. faecalis* (CCUG9997) | 13.8 | 33.2 | 0 | 1.09981817 | 19.4 |
| *E.coli* (NCTC13441) | 13.09 | 22.79 | 0.01527525 | 0.05 | 9.7 |
| *K. pneumoniae* (CCUG225T) | 30.6 | 38.49 | 0.42426407 | 1.48492424 | 7.89 |
| *P. aeruginosa* (CCUG17619) | 14.6 | 20.66 | 0 | 0.04582576 | 6.06 |
| *S. aureus* (CCUG17621) | 14 | 15.5166667 | 0 | 0.02516611 | 1.52 |

**S6 Table: qPCR ΔCT values between pure isolates and 24 hours**

| Species | Mean_Ct_Isolates | Mean_Ct_24 | SD_0 | SD_24 | ΔCt |
|---|---|---|---|---|---|
| *A. baumannii* (CCUG 19096T) | 13.63 | 13.22 | 0.01 | 0.03464102 | 0.41 |
| *E. faecalis* (CCUG9997) | 13.8 | 13.65 | 0 | 0.05 | 0.15 |
| *Ecoli* (NCTC13441) | 13.0866667 | 13.3066667 | 0.01527525 | 0.01154701 | 0.22 |
| *K. pneumoniae* (CCUG225T) | 30.6 | 25.8833333 | 0.42426407 | 0.1040833 | 4.72 |
| *P. aeruginosa* (CCUG17619) | 14.6 | 14.9866667 | 0 | 0.08082904 | 0.39 |
| *S. aureus* (CCUG17621) | 14 | 14.43 | 0 | 0.05196152 | 0.43 |

**S7 Table: Data retrieved from Oxford nanopore for evaluation of target species detection**

| Species | Total reads | Mean Read length | No. of reads aligned | Coverage | *E value* |
|---|---|---|---|---|---|
| *E. coli* (NCTC13441) | 160057 | 1745.7 | 7 | 0.33 | + |
| *P. aeruginosa* (CCUG17619) | 4210 | 1999.7 | 9 | 0.2 | + |
| *K. pneumoniae* (CCUG225T) | 59953 | 1708.5 | 425 | 13 | + |
| *A. baumannii* (CCUG 19096T) | 49318 | 1869.3 | 49 | 2 | + |
| *S. aureus* (CCUG17621) | 45358 | 1916.2 | 4677 | 95 | + |
| *E. faecalis* (CCUG9997) | 15298 | 1266.3 | NA* | NA* | NA* |

*NA, not available; +, highly significant*
